# Astrocytic morphology in the Medial Habenula: sex differences and modulatory factors

**DOI:** 10.64898/2026.06.29.735177

**Authors:** Cathaysa Rodríguez-Cedrés, Lucía Sangroniz-Beltrán, Nahia López, Naroe Delgado-Martín, Goiane Andueza-Peral, Paula Múgica-Susaeta, Eva Ducourneau, Paula Ospital, Sandra Beriain, María Ceprian, Jon Egaña-Huguet, Joaquín Piriz, Guillaume Ferreira, Susana Mato, Edgar Soria-Gómez

**Affiliations:** UPV/EHU–Achucarro, Leioa, Spain; UPV/EHU–Bilbao School of Engineering, Bilbao, Spain; NutriNeuro Lab, Université de Bordeaux, France; Ikerbasque, Bilbao, Spain; INSERM U1215 Neurocentre Magendie, Bordeaux, France; IIS Biobizkaia, Barakaldo, Spain

**Keywords:** Medial Habenula, Astrocytes, Sexual dimorphism, Inflammation

## Abstract

The medial habenula (MHb) is an epithalamic structure involved in aversive processing and emotional regulation, notable for its marked cellular heterogeneity and high astrocyte density. This cellular composition suggests that astrocytes may play an important role in MHb structure and plasticity, potentially contributing to the regulation of emotional states. <u>The aim of this study is to characterize</u> <u>sex-dependent astrocytic morphology in the MHb and determine how it is modulated by peripheral</u> <u>alterations and direct central manipulations</u>. A high-fat diet (HFD) was used as a model of metabolic stress, and systemic lipopolysaccharide (LPS) administration was used to induce a peripheral inflammatory challenge. At the central level, a chemogenetic approach using Gi-DREADDs under the GFAP promoter allowed selective modulation of astrocytic intracellular signalling independently of peripheral influences. Preliminary results indicate sex-dependent morphological differences in MHb astrocytes across all these experimental conditions, supporting the idea that MHb astrocytes are sensitive to both peripheral and central disturbances and may represent a key cellular substrate linking body-brain interactions with emotional regulation.

## 1. INTRODUCTION

Anxiety and depression are the most prevalent emotional disorders worldwide, representing a significant health and economic burden (1,2). Despite decades of research, the development of effective treatments remains challenging due to the complexity of the biological mechanisms involved. Increasing evidence suggests that emotional dysregulation is closely linked to neuroinflammatory processes and that metabolic and immune stressors can exacerbate neuroinflammation, thereby increasing vulnerability to emotional disorders (3,4).

Beyond classical limbic structures, the habenular complex has emerged as a key node in emotional regulation. This small but evolutionarily conserved epithalamic structure comprises two distinct subregions: the lateral habenula (LHb) and the medial habenula (MHb), which differ in their connectivity, cytoarchitecture, and molecular profiles (5). The LHb is the most extensively studied habenular subregion and has been strongly linked to the pathophysiology of depression and anxiety (6). In contrast, the MHb has received less attention, and its functions are not as clearly defined. However, it is believed to contribute to depression, anxiety, and fear, making it another critical area for the study of emotion and behavior. Furthermore, its neighboring vasculature exhibits increased transport function sensitive to peripheral modulators (including metabolic and immunological signals), suggesting that the MHb is particularly susceptible to peripheral influences on emotional pathophysiology (7).

Transcriptomic studies have revealed that approximately 50% of cells within the habenular complex are non-neuronal, with astrocytes representing a major population (5). Astrocytes perform a wide range of essential functions in the Central Nervous System (CNS): they regulate synaptic and inflammatory processes, maintain ionic balance, and participate in energy metabolism (8,9). When exposed to neurotoxic or inflammatory stimuli, astrocytes enter a reactive state characterized by cellular hypertrophy, increased branching complexity, and upregulation of markers such as GFAP and S100β (10, 11). These morphological changes have traditionally been interpreted as indicators of astrocyte activation (astrogliosis); however, it is now recognized that morphological analyses alone are insufficient to determine astrocyte functional state, as reactive astrocytes can adopt both neuroprotective and neurotoxic profiles (historically referred to as A2 and A1 states, respectively) depending on the nature of the stimulus and its duration (12). Moreover, external factors such as a high-fat diet (HFD) and lipopolysaccharide (LPS, a bacterial endotoxin that triggers a systemic immune response) have been shown to induce structural alterations in astrocytes and have been linked to the development of depressive and anxious symptoms, suggesting a connection between external disturbances and emotional dysfunction (3, 13, 14).

Despite the abundance of astrocytes in the MHb and their potential role in emotional regulation, their basal morphology and function remain largely uncharacterized. We hypothesize that MHb astrocytes undergo significant structural remodeling when exposed to metabolic, immune, or chemogenetic challenges, and that these morphological responses exhibit sexual dimorphism due to the well-known sex-dependent vulnerability to emotional and inflammatory stressors.

Here, we characterize MHb astrocyte morphology in male and female mice across three experimental conditions: metabolic stress (HFD), immune challenge (LPS), and chemogenetic modulation of astrocyte activity (using *Designer Receptors Exclusively Activated by Designer Drugs*: Gi-DREADDs). To capture morphology from complementary perspectives, we combine three approaches: skeleton-based analysis in Fiji, single-cell reconstruction (Simple Neurite Tracer, SNT), and Sholl analysis. As a first step, hippocampal (HC) astrocytes were included as a reference region for the DREADDs model, and astrocyte density was assessed by S100β quantification in the LPS group. Together, these analyses provide an initial characterization of MHb astrocyte morphology that we aim to extend across all conditions in future work.

## 2. MATERIALS AND METHODS

### 2.1. Animals and tissue preparation

C57BL/6 mice were used in every experimental group. Experiments were conducted in accordance with good laboratory practice and institutional guidelines for animal welfare, and all procedures were approved by the Animal Research Ethics Committee of the University of the Basque Country (UPV/EHU) (refs. M20/2019/199 and M30/2019/301). Experiments were approved the Region Aquitaine Veterinary Services (Direction Départementale de la Protection des Animaux, approval ID: B33-063-920) and by the local ethical committee (CEEA50) and the French Ministry of Agriculture and Forestry (authorization number APAFIS #15803).

#### High-fat diet

Brain tissue was provided by the laboratory of Dr. Guillaume Ferreira (University of Bordeaux, France). A total of 12 mice (6 males and 6 females) were provided with standard control diet (CD; A04 SAFE, Augy, France; offering 3.3 kcal/g consisting of 60% carbohydrate mostly from starch, and 3% fat), while another 12 mice (6 males and 6 females) received a high-fat high-sugar diet (HFD, D12451, Research Diets, New Brunswick, NJ; offering 4.7 kcal/g consisting of 24% fat (45% kcal), mostly saturated fat from lard, and 41% carbohydrate (35% kcal), with half coming from sucrose) (15). Mice had *ad libitum* access to water and food (either CD or HFD) for 12 weeks.

#### Lipopolysaccharide

Mice were randomly assigned to two groups. A total of 5 mice (2 males and 3 females) received an intraperitoneal injection of saline solution (SAL) as a control, while another 4 mice (1 male and 3 females) received an intraperitoneal injection of LPS (0.38 mg/kg; *Escherichia coli*, O55:B5, L6529, Sigma, dissolved in sterile saline).

#### Chemogenetics

A total of 8 mice (4 males and 4 females) received a bilateral infusion of an adeno-associated virus or AAV (ssAAV-8/2-hGFAP-hM4D(Gi)_mCherry-WPRE-hGHp(A)), which carries the Hm4Di receptor (Human Muscarinic Receiver 4). The Hm4Di receptor is a type of inhibitory DREADDs coupled to Gi proteins (16), whose expression is controlled by the astrocyte-specific GFAP promoter. Injections were performed at a 10-degree angle using a RWD stereotaxic apparatus (coordinates in mm: AP -1.55, ML ± 0.5, DV -3.3 and -3.1), injecting 100 nL of virus per injection with a microinjector (Nanoject III). The mice in the CTL group (n = 5, 3 males and 2 females) were injected with a control AAV carrying a mCherry marker (ssAAV-9/2-hGFAP-mCherry-WPRE-hGHp(A)), using the same surgical procedure. Before finishing the surgery, Meloxicam (2 mg/Kg) was administered subcutaneously. Surgical wounds were cleaned, sutured, and treated with topical lidocaine. All instruments and surfaces were disinfected before every surgery. After a 4-5-week recovery period, mice underwent behavioral testing, including a Fear Conditioning Test (data not shown). For this test, a single intraperitoneal injection of clozapine-N-oxide (CNO; 3 mg/kg) was administered 30 minutes before the acquisition phase. Following behavioral testing, mice were deeply anaesthetized with chloral hydrate (0.4 g/kg i.p., 4% in 0.1M PB) and transcardially perfused with 0.1M PB followed by 4% paraformaldehyde (PFA) in 0.1M PB. Brains were post-fixed in 4% PFA at 4°C for 24 hours and stored in 0.1M PB at 4°C until sectioning. Coronal sections (50 µm) were obtained using a Leica VT1000S vibratome and collected in 0.1M PB with 0.05% sodium azide.

### 2.2. Immunofluorescence

Immunofluorescence was performed on free-floating brain sections. Astrocytes were identified using two complementary markers: anti-GFAP (Alexa Fluor 488 α Mo; 1:500, Thermo Fisher Scientific) and anti-S100β ([EP1576Y], anti-rabbit; 1:500, Abcam, ab52642). Sections were coverslipped using Fluoroshield Mounting Medium with DAPI (F6057; Sigma-Aldrich; St. Louis, USA), sealed with nail polish, and slices were stored at 4°C.

### 2.3. Image acquisition

Fluorescence images were acquired with a ZEISS Apotome 3 fluorescence microscope (SGIker, UPV/EHU) using a 20X/0.8 NA air objective and ZEN 2.6 Pro software.

### 2.4. Morphological analysis

Astrocyte morphology was first analyzed with Fiji (ImageJ 2.14.0) using a skeleton-based approach. GFAP images were split from the original channels, and Z-stacks were projected using maximum intensity. Brightness and contrast were automatically adjusted, greyscale LUTs were applied, and an unsharp mask was used to enhance structural edges.

Five astrocytes per MHb were selected using the Region of Interest (ROI) tool (left and right hemispheres), for a total of 10 astrocytes per slice (with 3-5 slices analyzed per animal, depending on tissue availability and quality). These regions were duplicated, converted to 8-bit, and arranged to create binary masks, which were manually refined to delineate cell boundaries. The binarized images were then skeletonized and analyzed, extracting the following parameters (Figure 1, A):

- Number of branches: total number of extensions from the cell body.
- Mean branch length (μm): average length of all individual branches.
- Maximum branch length (μm): length of the single longest branch.
- Longest-shortest path length (μm): the longest of all shortest paths between any two points within the astrocyte skeleton.

**Figure 1.**
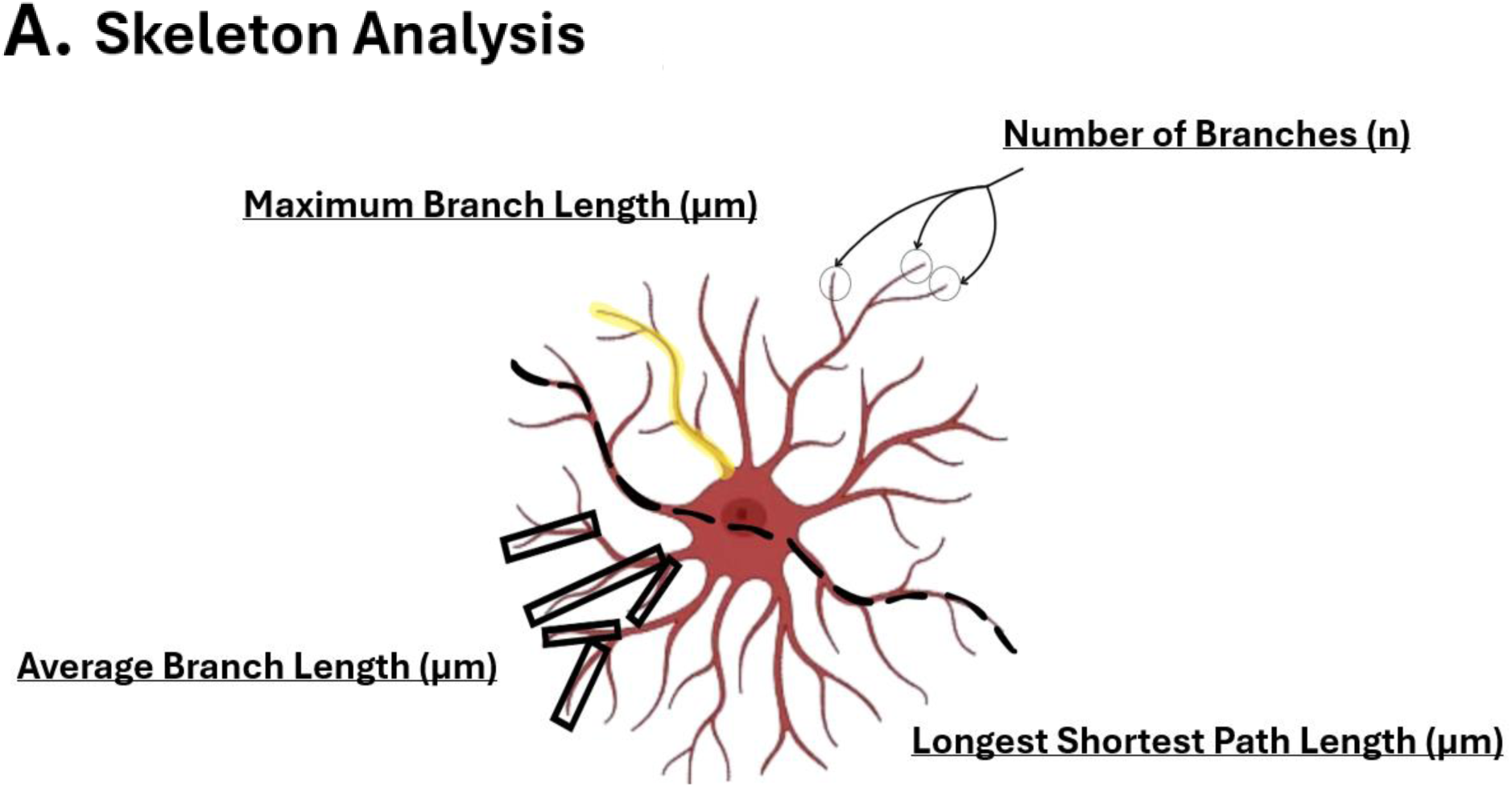
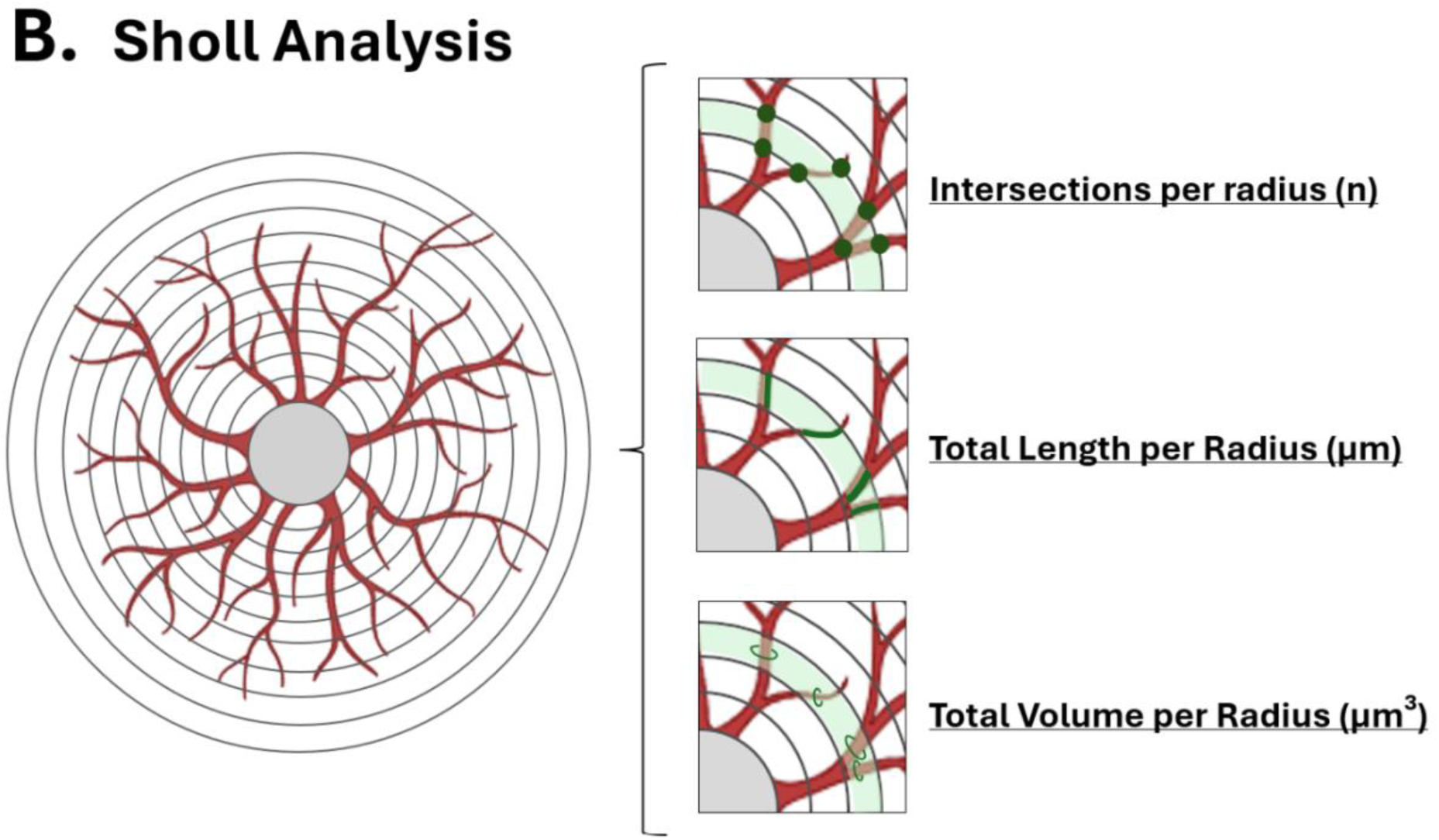
Morphological parameters measured in MHb astrocytes. **(A)** Skeleton analysis. Astrocytes were skeletonized in Fiji to extract four morphological parameters: number of branches (n), maximum branch length (µm), average branch length (µm) and longest shortest path length (µm), defined as the longest of all shortest paths between any two points of the skeleton. **(B)** Sholl analysis. Concentric circles centred on the soma at fixed radial intervals were used to quantify branching complexity as a function of distance from the cell body, measuring the number of intersections (n), total process length (µm) and total process volume (µm³) within each radius.

This skeleton-based analysis was performed on MHb astrocytes across all experimental groups. To provide a reference framework, it was additionally applied to hippocampal (CA1) astrocytes in the DREADDs model, allowing a comparative assessment of structural complexity between brain regions.

Astrocyte morphology was further characterized using the SNT (Simple Neurite Tracer) plugin in Fiji, applied to the MHb of the DREADDs group. Unlike the skeleton-based method, SNT works directly on the complete Z-stack rather than on a Z-projection, preserving depth information. Individual astrocytes were reconstructed by manually tracing their processes from the soma through the primary and secondary branches, establishing the structural hierarchy of each cell. From these traces, the same parameters described above were obtained, together with volume-related measurements (cell volume and branch volume), providing a more detailed, three-dimensional description of astrocyte morphology.

Finally, branching complexity was assessed by Sholl analysis on the same SNT reconstructions. Concentric circles were placed at fixed radial intervals (4 μm) from the soma, and within each circle, the number of intersections, total process length, and estimated volume were quantified (Figure 1, B). This allowed us to describe how astrocytic complexity varies with distance from the soma. As with SNT, this analysis was performed on the MHb of the DREADDs group.

All three analyses are intended to be extended to the remaining experimental groups in future work.

### 2.5. Quantification

A programmatic method was developed to quantify astrocytes objectively, replacing manual annotation, which is time-consuming and subjective. The workflow was designed as a human-in-the-loop pipeline, where automated detection is supervised and validated by the researcher’s expertise.

The pipeline was implemented in Python using OpenCV. Images are first converted to grayscale and inverted, then binarized with a local adaptive threshold to compensate for uneven illumination. A Gaussian blur removes noise while preserving cellular detail, and a final fixed threshold produces a clean binary mask.

The mask is refined through a sequence of erosion and dilation operations. An initial erosion separates cells in near contact; an iterated dilation closes the internal holes within each astrocyte; and a final erosion restores the original object size. The dilation step is essential since quantification is based on counting contours; unclosed holes would otherwise be detected as separate objects, inflating the count. Objects are then detected as a flat list of contours.

To prevent unwanted objects, each contour passes through a series of inclusion filters based on area, perimeter, and circularity. The area bounds discard punctual noise and excessively large artifacts. Near-zero-perimeter contours are removed as degenerate detections; and, because astrocytes are stellate, objects approaching a perfect circle are treated as noise.

The final count is obtained by removing duplicates, the centroid of each valid contour is computed from image moments, and centroids lying within a minimum distance threshold of an already accepted centroid are discarded, assuring a single cell cannot be counted more than once (Figure 2).

**Figure 2.**
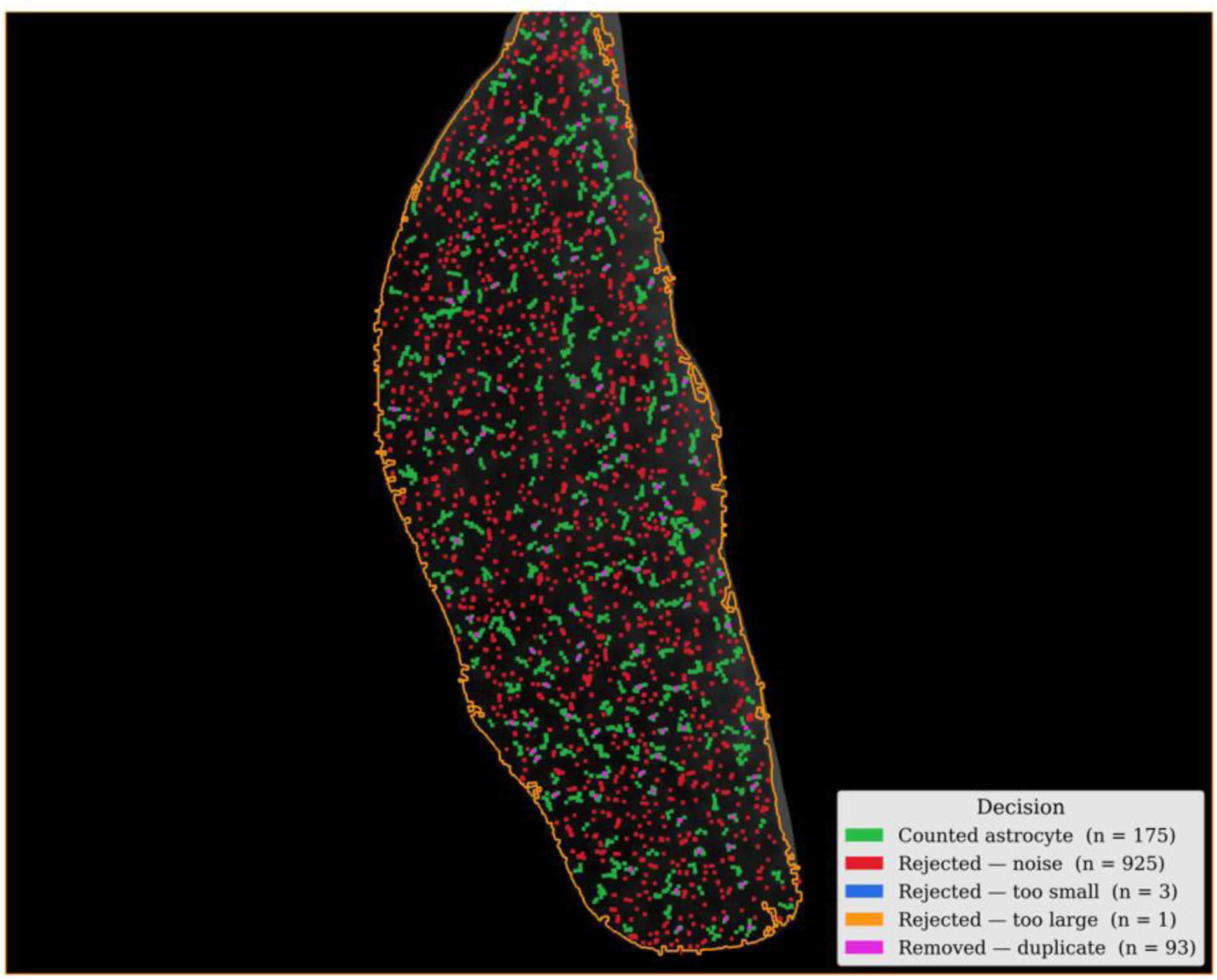
Algorithmic decision map for automated astrocyte quantification in the MHb. Representative coronal section of the medial habenula (MHb) immunostained for S100β, with the analysed region delineated (orange outline). Every detected object is colour-coded by the decision taken by the algorithm: counted astrocyte (green), rejected as noise (red), rejected by area (too small, blue; too large, orange), and removed as a duplicate (magenta). Object counts for each category are indicated in the legend. This single view illustrates what the algorithm keeps and discards.

To assess reliability, confidence scores are added. The per-object confidence score combines the local contrast with its surroundings, a plausible-area criterion, and its circularity to be within the expected range. A global, image-level confidence score is computed as a weighted combination of mean individual confidence, size consistency, object density, and the contour acceptance rate. These scores, together with automatic warnings, flag images that need manual review, closing the human-in-the-loop.

For each image, the output is written to a spreadsheet that reports the morphological variables along with the confidence variables. All processing parameters are centralized in a single configuration file to ensure full reproducibility.

## 3. RESULTS

### 3.1. High-fat diet

Regarding male mice (Figure 3), a HFD induced structural changes in MHb astrocytes. Analysis showed a significant increase in the number of branches in the HFD group compared to the CD group (Figure 3, D; *p < 0.05). No significant differences were observed in maximum branch length (Figure 3, E), average branch length (Figure 3, F), or the longest-shortest path length (Figure 3, G). These findings are supported by the skeletonized reconstructions, which clearly reveal a denser network with more branches in HFD astrocytes (Figure 3, C.2).

**Figure 3.**
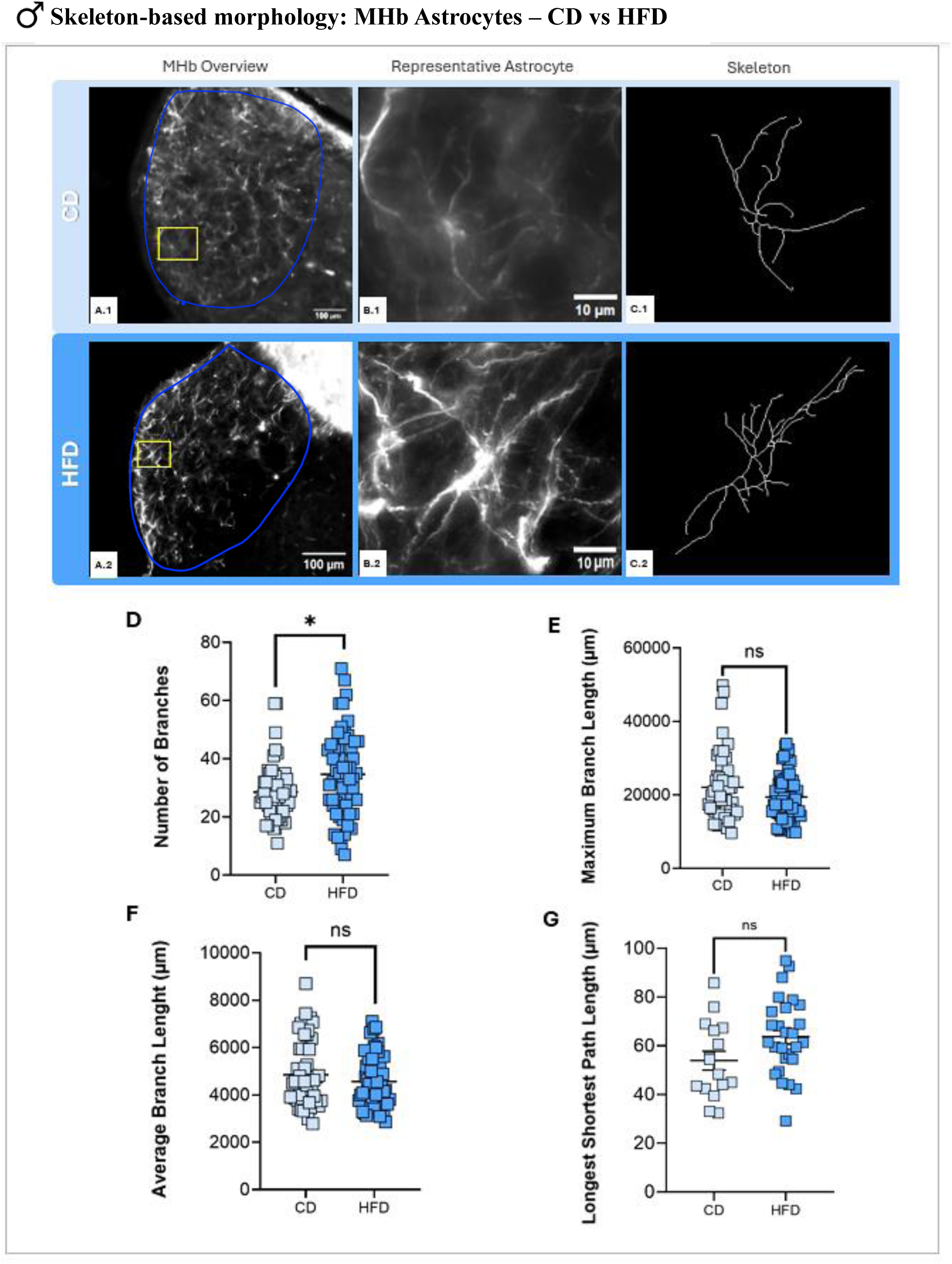
Manual astrocyte morphological analysis under metabolic stress (CD vs HFD) in the MHb of male mice. **A-C**: Representative images for control group (CD: A.1-C.1), and experimental group (HFD: A.2-C.2). MHb overview (A; scale bar, 100 µm), magnified representative astrocyte (B; scale bar, 10 µm) and corresponding skeleton (C). **D–G**: Skeleton-based morphological parameters quantified per astrocyte. **D**: number of branches, **E**: maximum branch length, **F**: average branch length, and **G**: longest shortest path length. A two-tailed unpaired Student’s T-test was used for comparisons. Normal distribution and equal variances were assumed. Values are expressed as mean ± SEM. Significance levels are indicated as follows: ns = not significant; *p < 0.05.

On the other hand, the diet did not trigger significant changes in the female cohort (Figure 4). No differences were detected between CD and HFD groups in the number of branches (Figure 4, D), maximum branch length (Figure 4, E), average branch length (Figure 4, F), or the longest-shortest path length (Figure 4, G). However, the skeletonized images still show a slightly more complex pattern with more branches in HFD females compared to controls (Figure 4, C.2).

**Figure 4.**
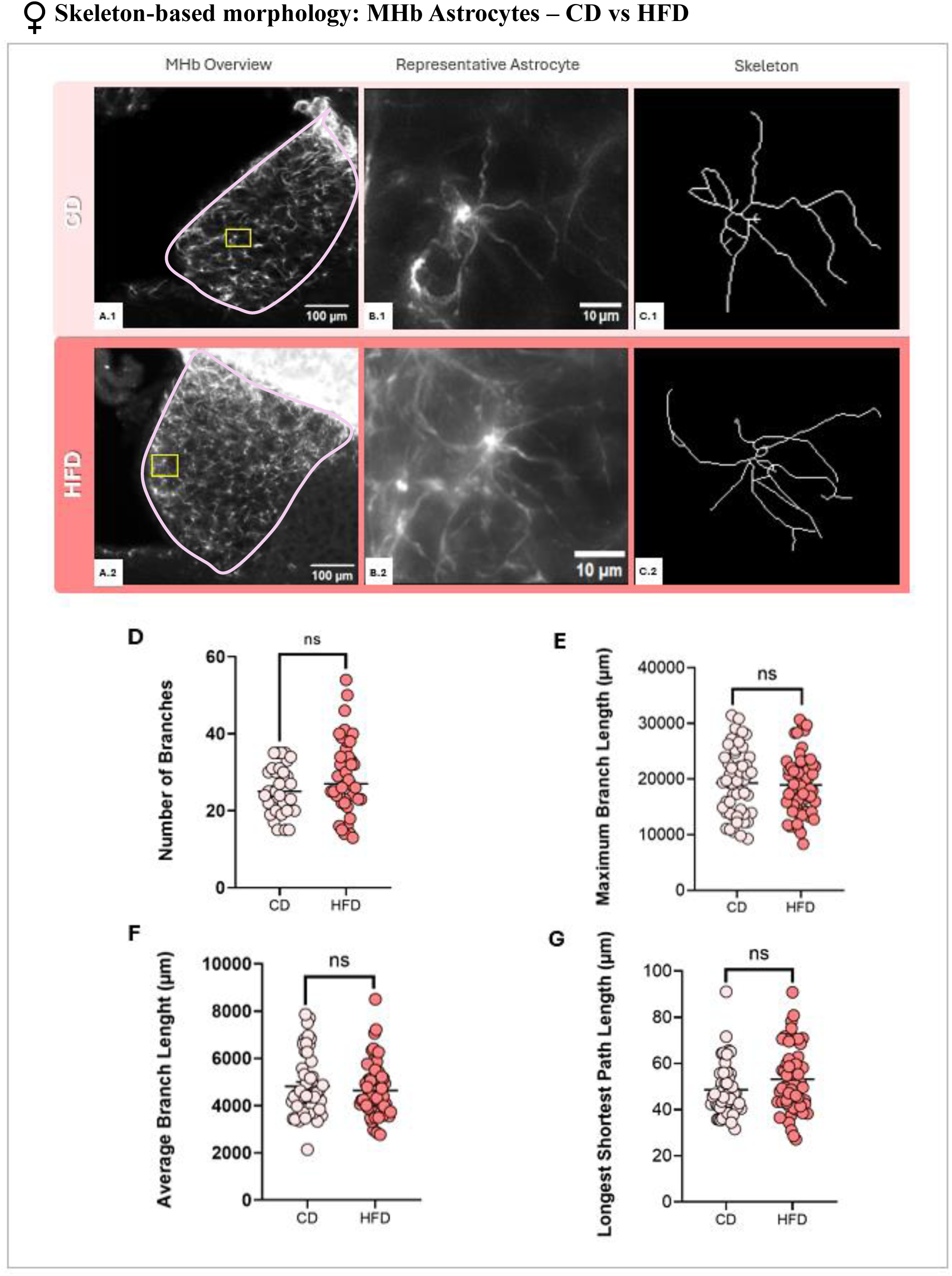
Manual astrocyte morphological analysis under metabolic stress (CD vs HFD) in the MHb of female mice. **A-C**: Representative images for control group (CD: A.1-C.1), and experimental group (HFD: A.2-C.2). MHb overview (A; scale bar, 100 µm), magnified representative astrocyte (B; scale bar, 10 µm) and corresponding skeleton (C). **D–G**: Skeleton-based morphological parameters quantified per astrocyte. **D**: number of branches, **E**: maximum branch length, **F**: average branch length, and **G**: longest shortest path length. A two-tailed unpaired Student’s T-test was used for comparisons. Normal distribution and equal variances were assumed. Values are expressed as mean ± SEM. Significance levels are indicated as follows: ns = not significant.

### 3.2. Lipopolysaccharide

As shown in Figure 5, male mice exhibited a subtle decrease in MHb astrocyte complexity following systemic LPS administration. Analysis revealed a significant reduction only in the number of branches in the LPS group compared to CTL (Figure 5, D; *p < 0.05). No significant differences were found in maximum branch length (Figure 5, E), average branch length (Figure 5, F), or the longest-shortest path length (Figure 5, G). This reduction is visually reflected in the skeletonized reconstructions, showing fewer processes following treatment (Figure 5, C.2).

**Figure 5.**
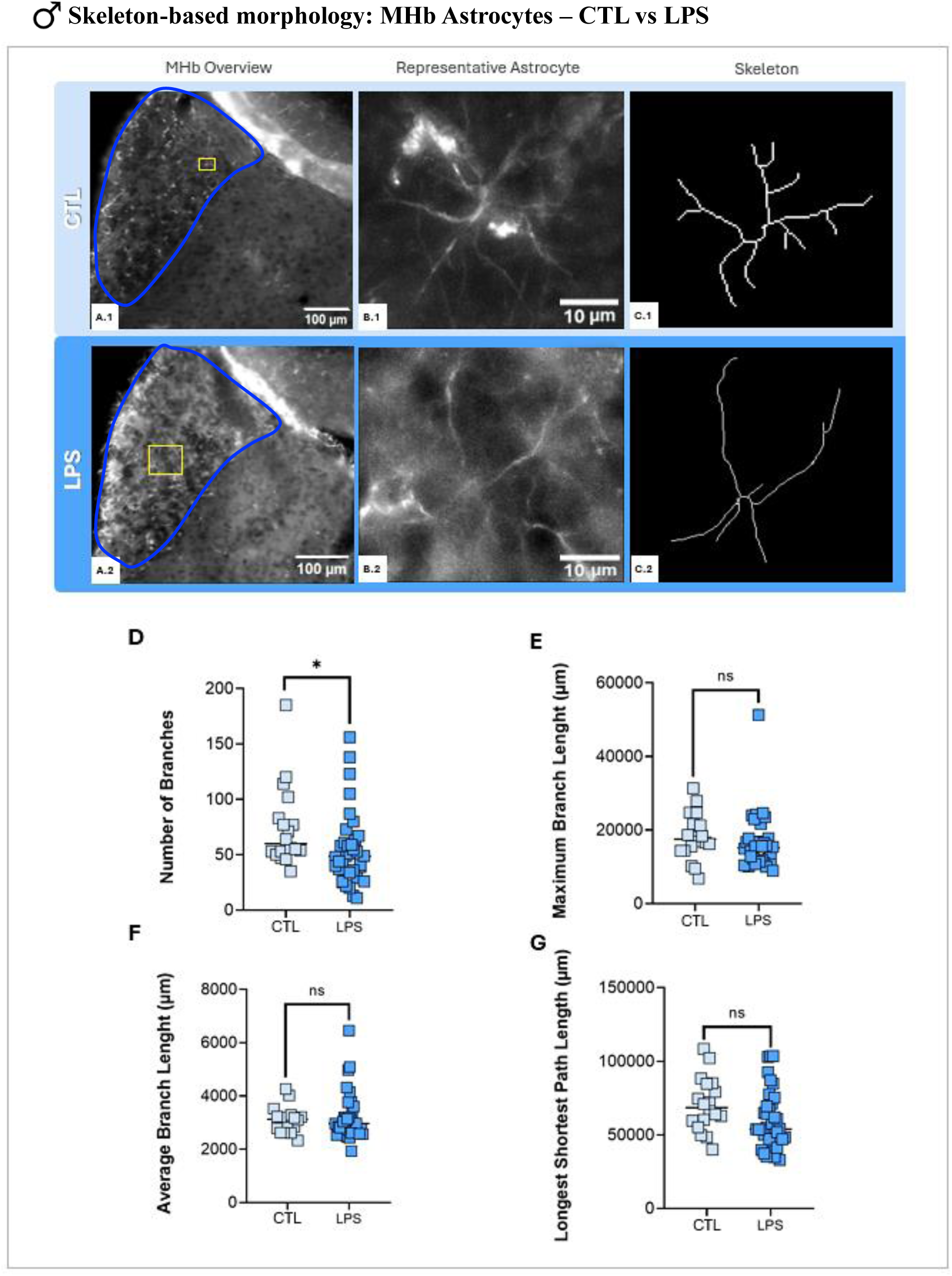
Manual astrocyte morphological analysis under immune challenge (CTL vs LPS) in the MHb of male mice. **A-C**: Representative images for control group (CTL: A.1-C.1), and experimental group (LPS: A.2-C.2). MHb overview (A; scale bar, 100 µm), magnified representative astrocyte (B; scale bar, 10 µm) and corresponding skeleton (C). **D–G**: Skeleton-based morphological parameters quantified per astrocyte. **D**: number of branches, **E**: maximum branch length, **F**: average branch length, and **G**: longest shortest path length. A two-tailed unpaired Student’s T-test was used for comparisons. Normal distribution and equal variances were assumed. Values are expressed as mean ± SEM. Significance levels are indicated as follows: ns = not significant; *p < 0.05.

In contrast, the morphological changes were considerably more pronounced and statistically significant in the female cohort (Figure 6). LPS-treated females exhibited a highly significant decrease in both the number of branches (Figure 6, D; ***p < 0.001) and the longest-shortest path length (Figure 6, G; ***p < 0.001). Additionally, a significant reduction was observed in the maximum branch length (Figure 6, E; **p < 0.01), whereas the average branch length remained unaffected (Figure 6, F). These results are visually supported by the skeletonized images, which show a clear loss of complexity and fewer branches in LPS females (Figure 6, C.2).

**Figure 6.**
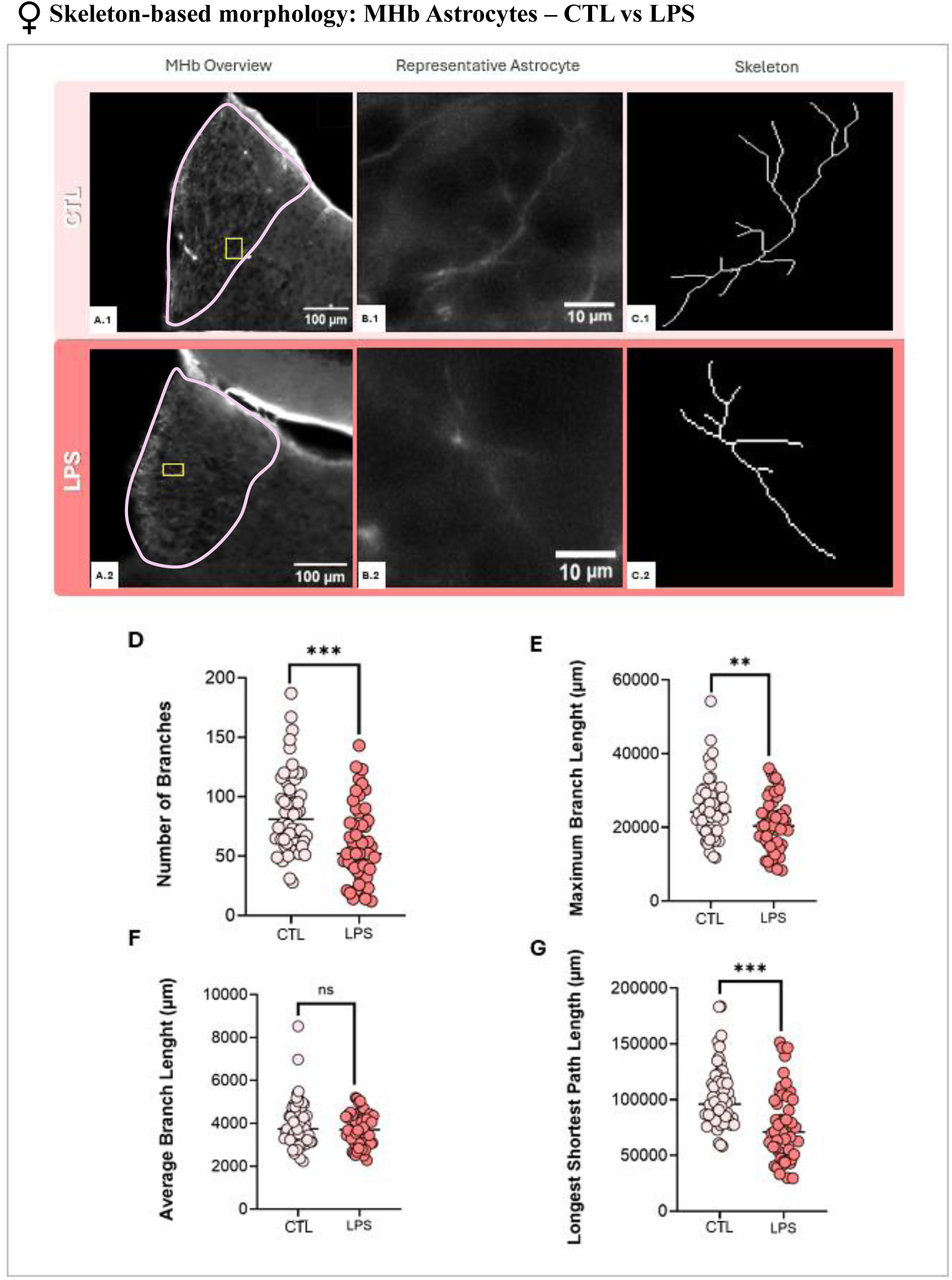
Manual astrocyte morphological analysis under immune challenge (CTL vs LPS) in the MHb of female mice. **A-C**: Representative images for control group (CTL: A.1-C.1), and experimental group (LPS: A.2-C.2). MHb overview (A; scale bar, 100 µm), magnified representative astrocyte (B; scale bar, 10 µm) and corresponding skeleton (C). **D–G**: Skeleton-based morphological parameters quantified per astrocyte. **D**: number of branches, **E**: maximum branch length, **F**: average branch length, and **G**: longest shortest path length. A two-tailed unpaired Student’s T-test was used for comparisons. Normal distribution and equal variances were assumed. Values are expressed as mean ± SEM. Significance levels are indicated as follows: ns = not significant; **p < 0.01, ***p < 0.001.

Male mice showed no statistically significant differences in either parameter (Figure 7). However, a slight upward trend is visible in the LPS group, which tended to show a higher number of detected s100bβ + astrocytes (Figure 7, A).

**Figure 7.**
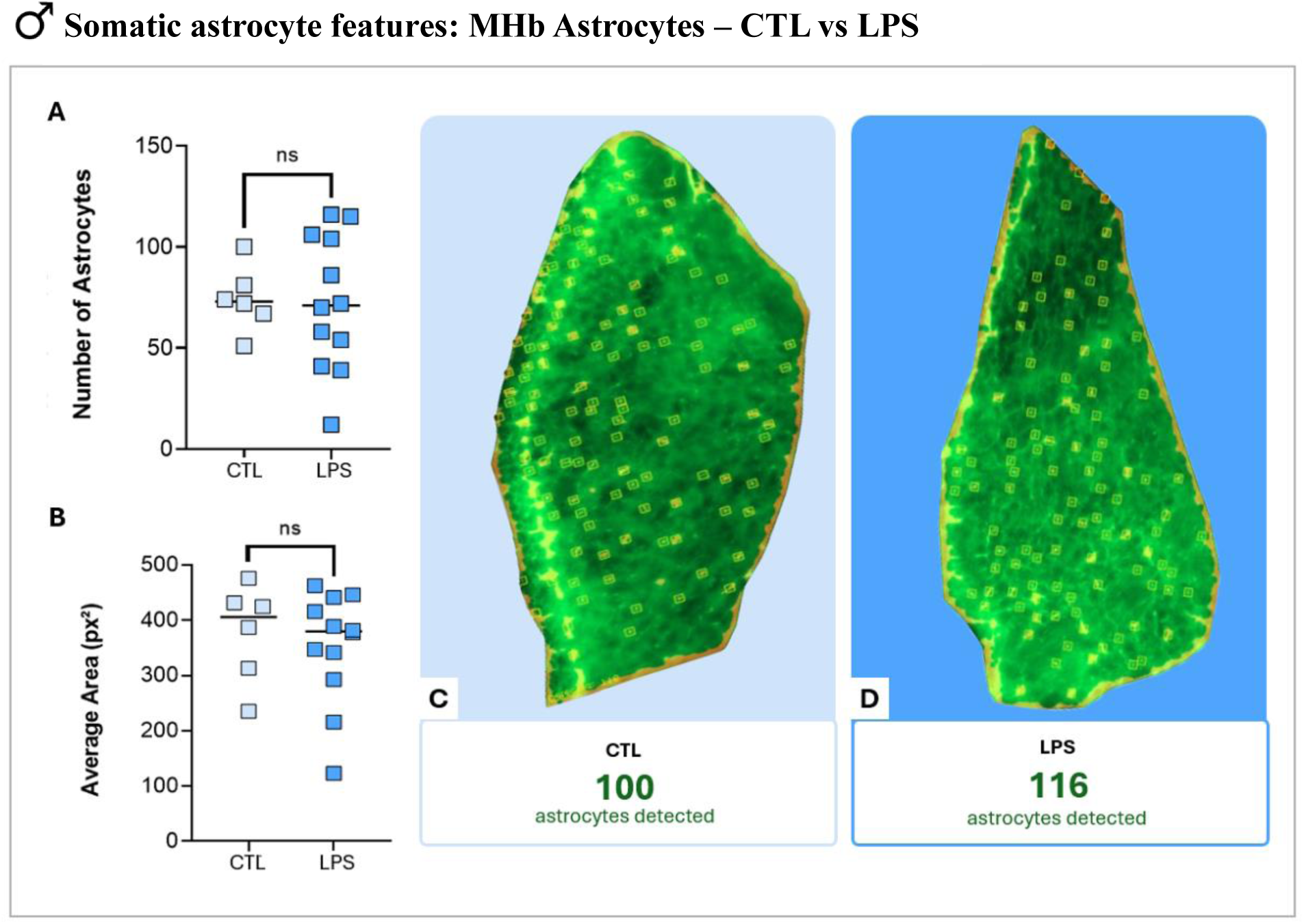
S100β+ astrocyte density and area analysis under LPS-induced inflammation (CTL vs LPS) in the MHb of male mice. **A**: total number of astrocytes, and **B**: average astrocyte area measured in square pixels (px²). **C–D**: Representative overview masks of the MHb showing automated cell detection for the control group (CTL; C) and the experimental group (LPS; D), with the respective number of detected astrocytes indicated below. A two-tailed unpaired Student’s T-test was used for comparisons. Normal distribution and equal variances were assumed. Values are expressed as mean ± SEM. Significance levels are indicated as follows: ns = not significant.

In contrast, female mice showed a statistically significant inflammatory response (Figure 8). Following LPS administration, the number of detected S100β+ astrocytes decreased compared to the CTL group (Figure 8, A; **p < 0.01). Regarding cell area, while not statistically significant, a slight increasing trend was observed in this variable in the LPS group (Figure 8, B).

**Figure 8.**
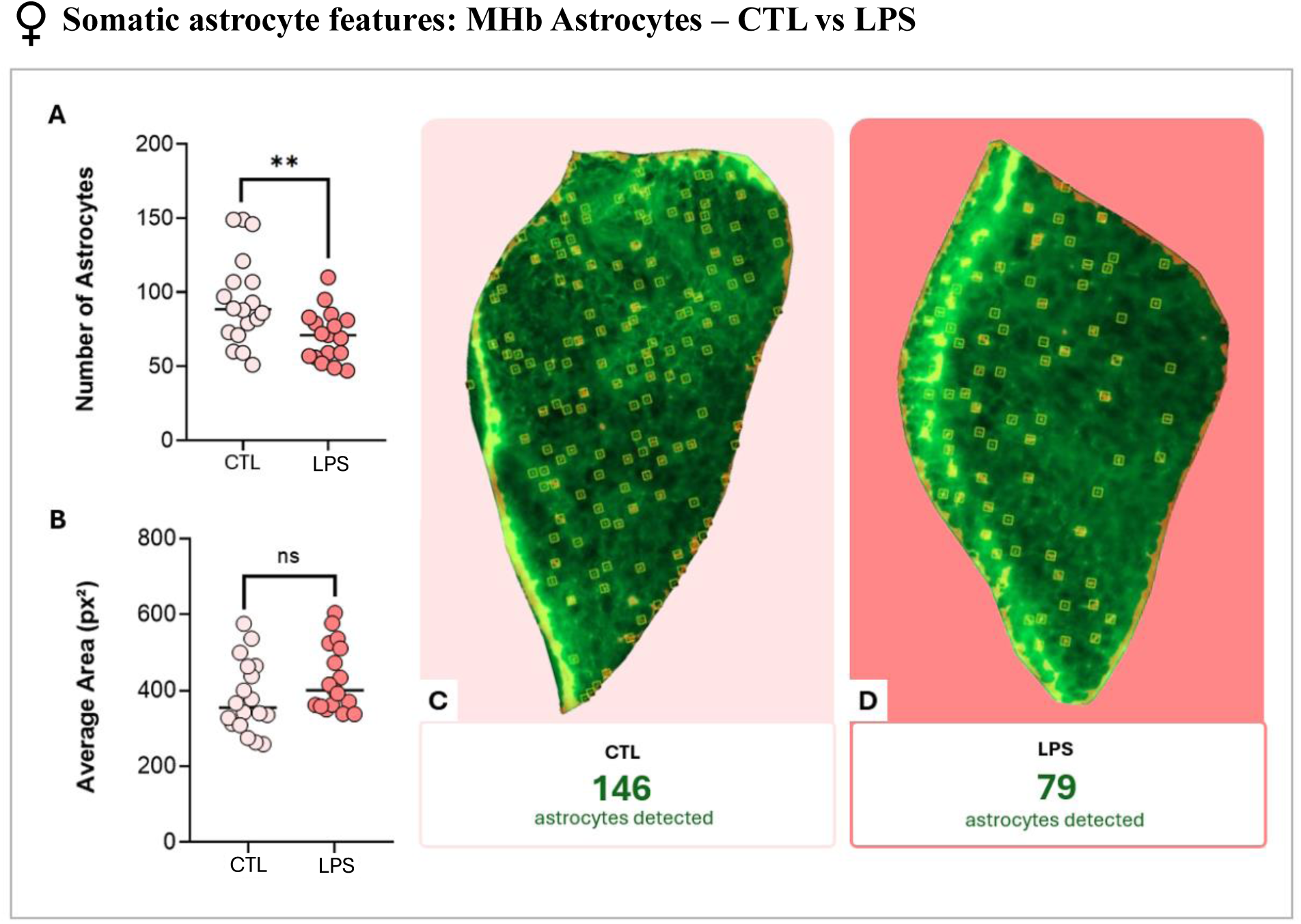
S100β+ astrocyte density and area analysis under LPS-induced inflammation (CTL vs LPS) in the MHb of female mice. **A**: total number of astrocytes, and **B**: average astrocyte area measured in square pixels (px²). **C–D**: Representative overview masks of the MHb showing automated cell detection for the control group (CTL; C) and the experimental group (LPS; D), with the respective number of detected astrocytes indicated below. A two-tailed unpaired Student’s T-test was used for comparisons. Normal distribution and equal variances were assumed. Values are expressed as mean ± SEM. Significance levels are indicated as follows: ns = not significant, **p < 0.01.

### 3.3. Chemogenetics

In male mice (Figure 9), DREADDs-expressing astrocytes showed a significant increase in the number of branches compared to CTL (Figure 9, D; ***p < 0.001), as well as a longer longest-shortest path length (Figure 9, G; **p < 0.01). No significant differences were found in maximum branch length (Figure 9, E) or average branch length (Figure 9, F). These results are visually supported by the skeletonized reconstructions, which reveal a denser and more complex branching pattern in DREADDs astrocytes (Figure 9, C.2).

**Figure 9.**
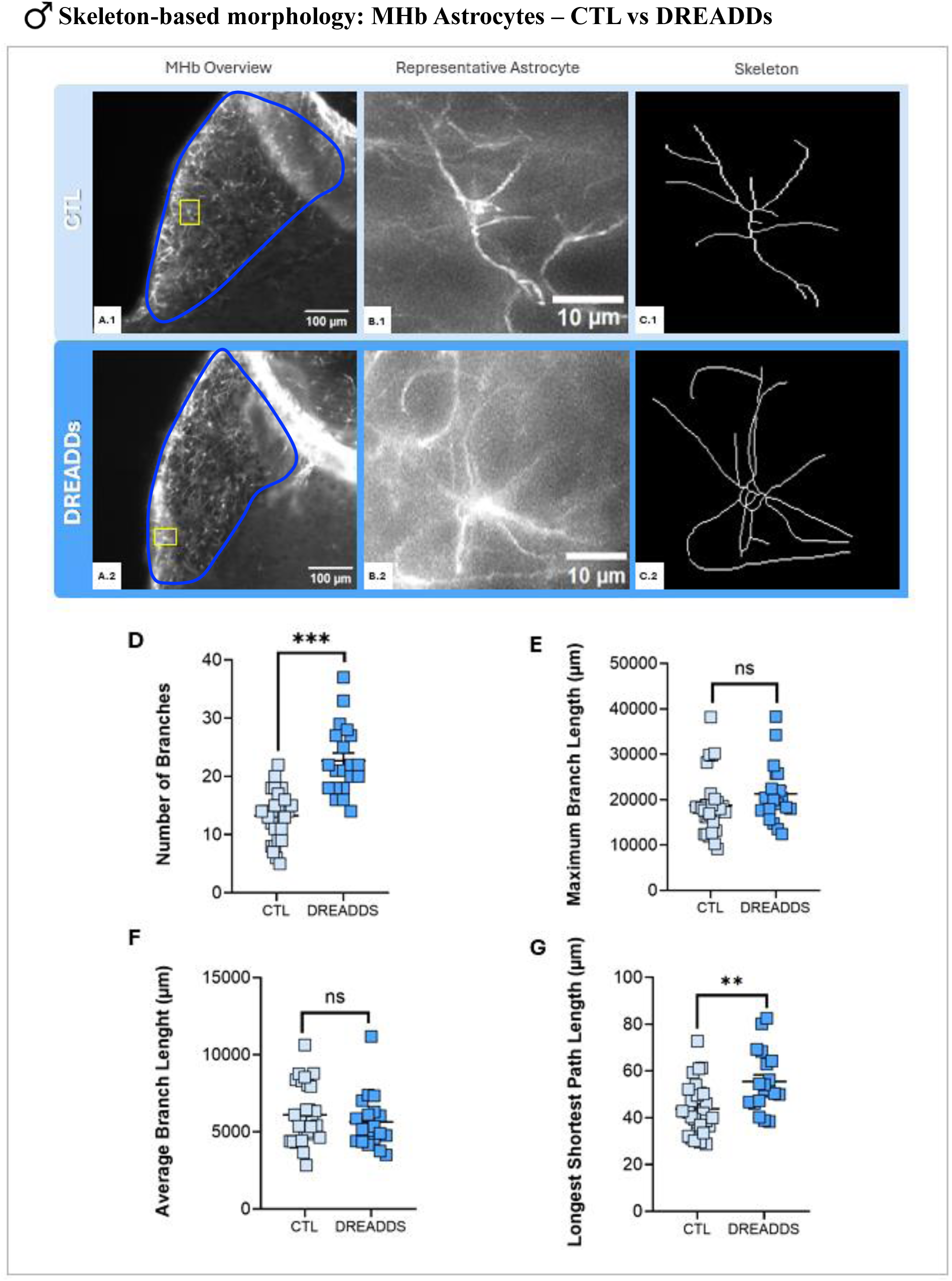
Manual astrocyte morphological analysis under chemogenetic manipulation (CTL vs DREADDs) in the MHb of male mice. **A-C**: Representative images for control group (CTL: A.1-C.1), and experimental group (DREADDs: A.2-C.2). MHb overview (A; scale bar, 100 µm), magnified representative astrocyte (B; scale bar, 10 µm) and corresponding skeleton (C). **D–G**: Skeleton-based morphological parameters quantified per astrocyte. **D**: number of branches, **E**: maximum branch length, **F**: average branch length, and **G**: longest shortest path length. A two-tailed unpaired Student’s T-test was used for comparisons. Normal distribution and equal variances were assumed. Values are expressed as mean ± SEM. Significance levels are indicated as follows: ns = not significant; **p < 0.01; ***p < 0.001. *Note:* a*ll animals in this cohort underwent a fear conditioning protocol, including the administration of CNO in the acquisition phase*.

In female mice (Figure 10), DREADDs expression led to a significant increase in both the number of branches (Figure 10, D; *p < 0.05) and the average branch length (Figure 10, F; *p < 0.05). In contrast, no significant changes were observed in maximum branch length (Figure 10, E) or the longest-shortest path length (Figure 10, G). Visually, the skeletonized reconstructions confirm an increase in the complexity and expansion of branches in DREADDs astrocytes (Figure 10, C.2).

**Figure 10.**
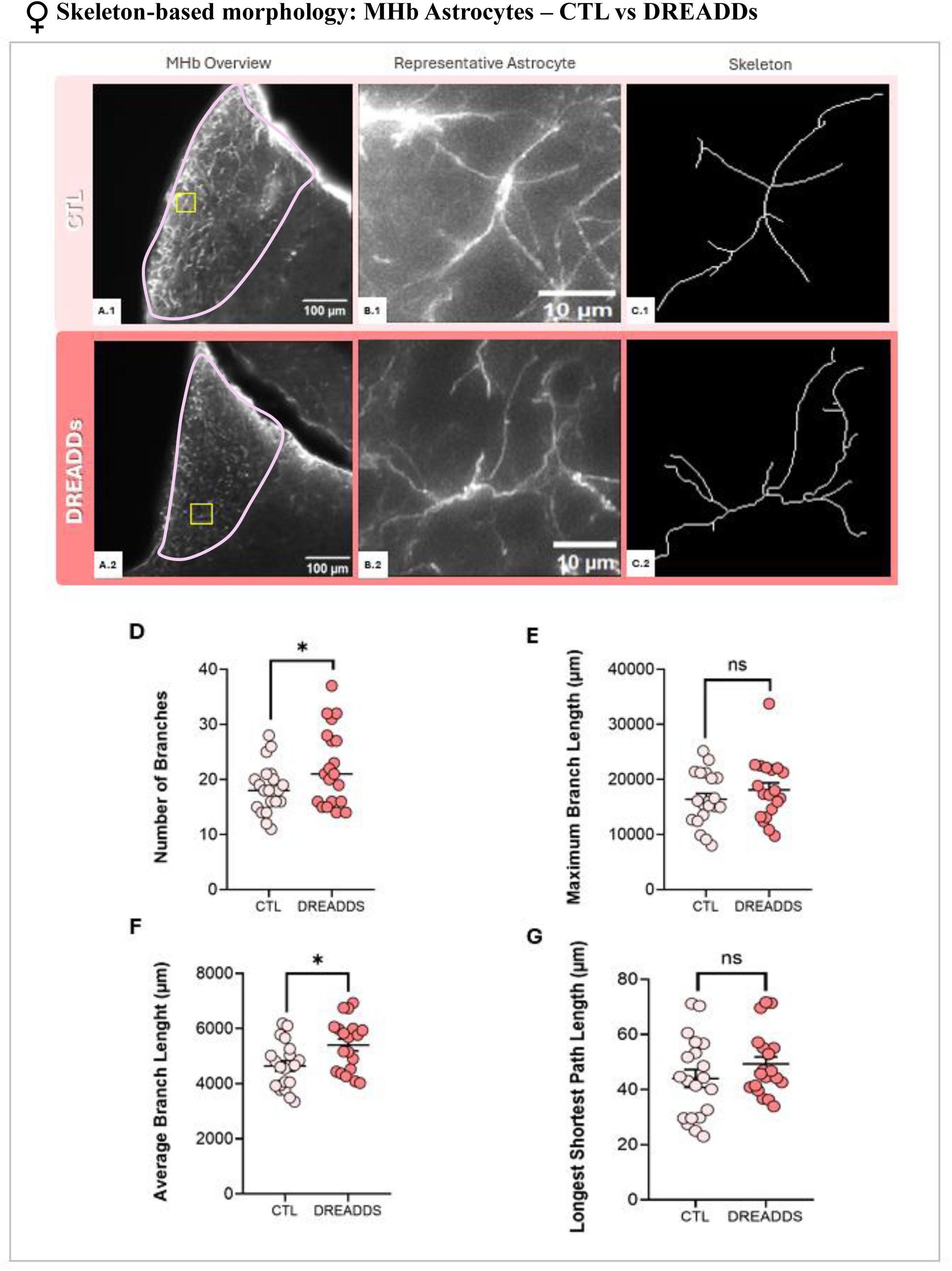
Manual astrocyte morphological analysis under chemogenetic manipulation (CTL vs DREADDs) in the MHb of female mice. **A-C**: Representative images for control group (CTL: A.1-C.1), and experimental group (DREADDs: A.2-C.2). MHb overview (A; scale bar, 100 µm), magnified representative astrocyte (B; scale bar, 10 µm) and corresponding skeleton (C). **D–G**: Skeleton-based morphological parameters quantified per astrocyte. **D**: number of branches, **E**: maximum branch length, **F**: average branch length, and **G**: longest shortest path length. A two-tailed unpaired Student’s T-test was used for comparisons. Normal distribution and equal variances were assumed. Values are expressed as mean ± SEM. Significance levels are indicated as follows: ns = not significant; *p < 0.05. *Note:* a*ll animals in this cohort underwent a fear conditioning protocol, including the administration of CNO in the acquisition phase*.

In male mice (Figure 11), chemogenetic manipulation did not induce any significant changes in CA1 astrocyte morphology. No statistically significant differences were observed between the CTL and DREADDs groups in the number of branches (Figure 11, D), maximum branch length (Figure 11, E), average branch length (Figure 11, F), or the longest-shortest path length (Figure 11, G).

**Figure 11.**
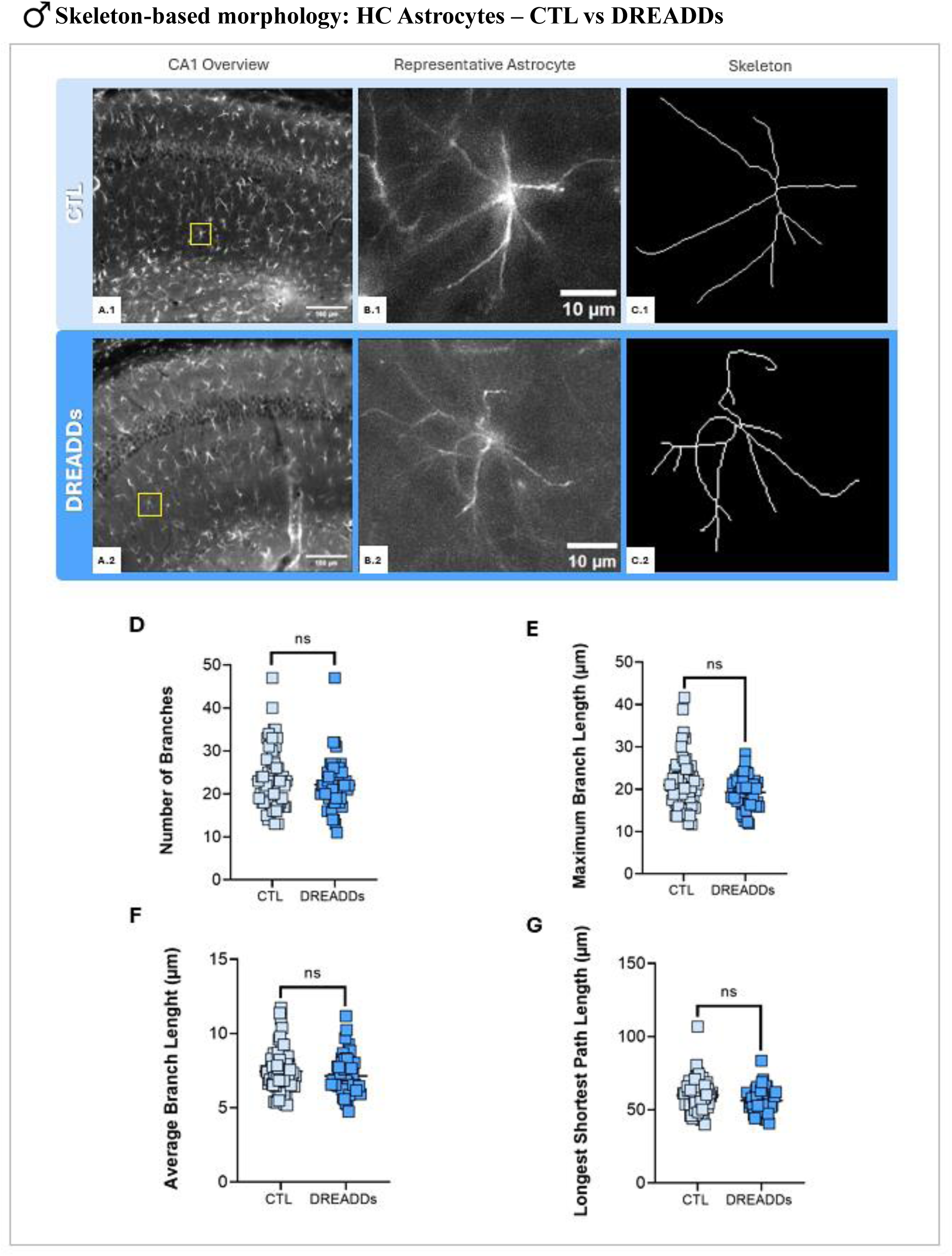
Manual HC-astrocyte morphological analysis after MHb-chemogenetic manipulation (CTL vs DREADDs) of male mice. **A-C**: Representative images for control group (CTL: A.1-C.1), and experimental group (DREADDs: A.2-C.2). CA1 overview (A; scale bar, 100 µm), magnified representative astrocyte (B; scale bar, 10 µm) and corresponding skeleton (C). **D–G**: Skeleton-based morphological parameters quantified per astrocyte. **D**: number of branches, **E**: maximum branch length, **F**: average branch length, and **G**: longest shortest path length. A two-tailed unpaired Student’s T-test was used for comparisons. Normal distribution and equal variances were assumed. Values are expressed as mean ± SEM. Significance levels are indicated as follows: ns = not significant. *Note:* a*ll animals in this cohort underwent a fear conditioning protocol, including the administration of CNO in the acquisition phase*.

In female mice (Figure 12), DREADDs expression led to distinct morphological alterations in the CA1 region. Astrocytes showed a significant increase in the number of branches compared to CTL (Figure 12, D; *p < 0.05). Conversely, a significant decrease was observed in the average branch length (Figure 12, F; *p < 0.05). No significant changes were found in maximum branch length (Figure 12, E) or the longest-shortest path length (Figure 12, G).

**Figure 12.**
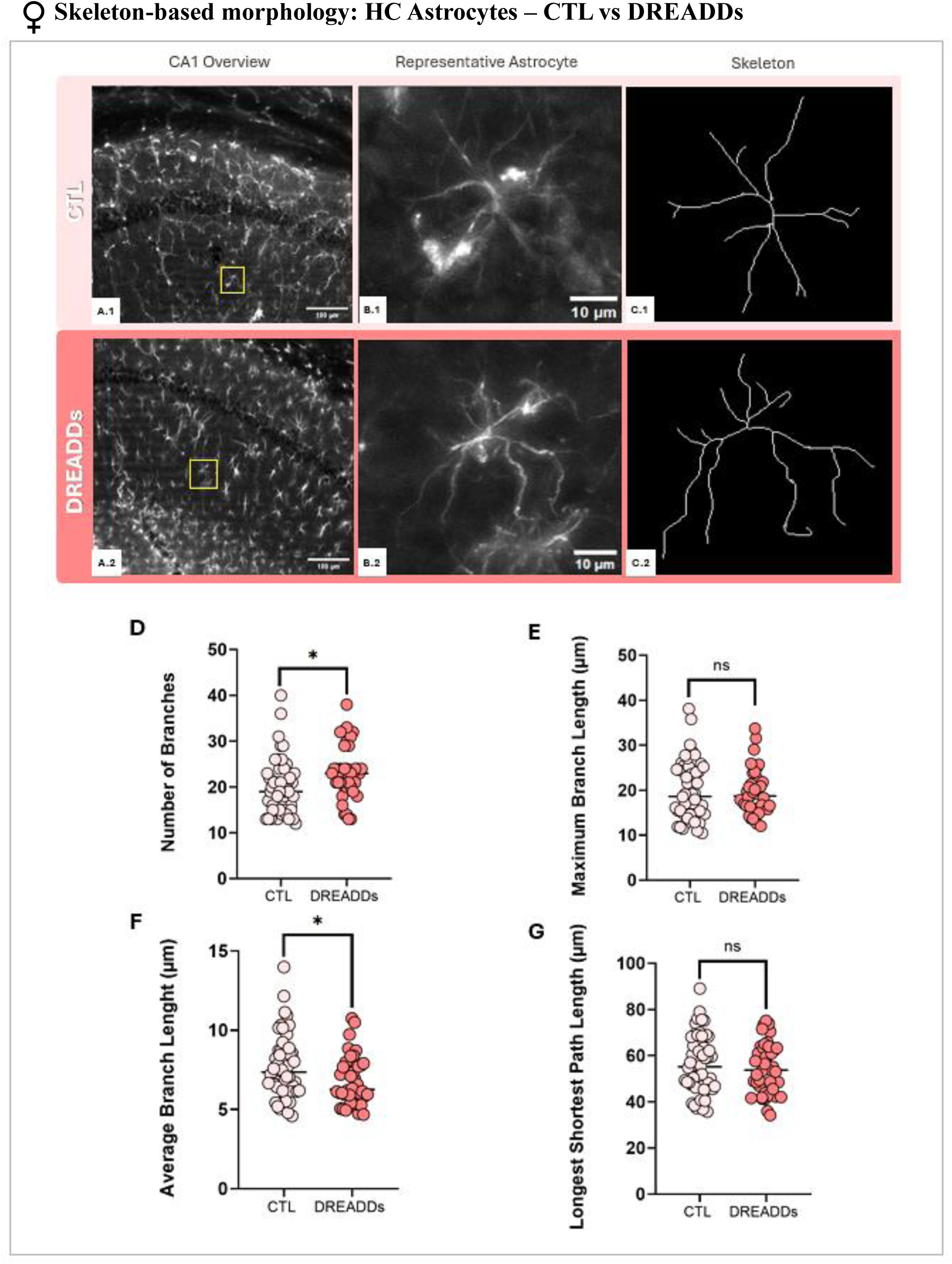
Manual HC-astrocyte morphological analysis after MHb-chemogenetic manipulation (CTL vs DREADDs) of female mice. **A-C**: Representative images for control group (CTL: A.1-C.1), and experimental group (DREADDs: A.2-C.2). CA1 overview (A; scale bar, 100 µm), magnified representative astrocyte (B; scale bar, 10 µm) and corresponding skeleton (C). **D–G**: Skeleton-based morphological parameters quantified per astrocyte. **D**: number of branches, **E**: maximum branch length, **F**: average branch length, and **G**: longest shortest path length. A two-tailed unpaired Student’s T-test was used for comparisons. Normal distribution and equal variances were assumed. Values are expressed as mean ± SEM. Significance levels are indicated as follows: ns = not significant, *p < 0.05. *Note:* a*ll animals in this cohort underwent a fear conditioning protocol, including the administration of CNO in the acquisition phase*.

From a more detailed morphological perspective using SNT and Sholl analysis, Figure 13 shows that chemogenetic manipulation in male mice did not lead to statistically significant changes in any of the individual parameters measured by SNT (Figure 13, D–I). However, in the DREADD group, a non-significant trend toward an increase in the average branch volume and total astrocyte volume was observed. This slight increase was consistently reflected in the Sholl analysis profiles (Figure 13, L).

**Figure 13.**
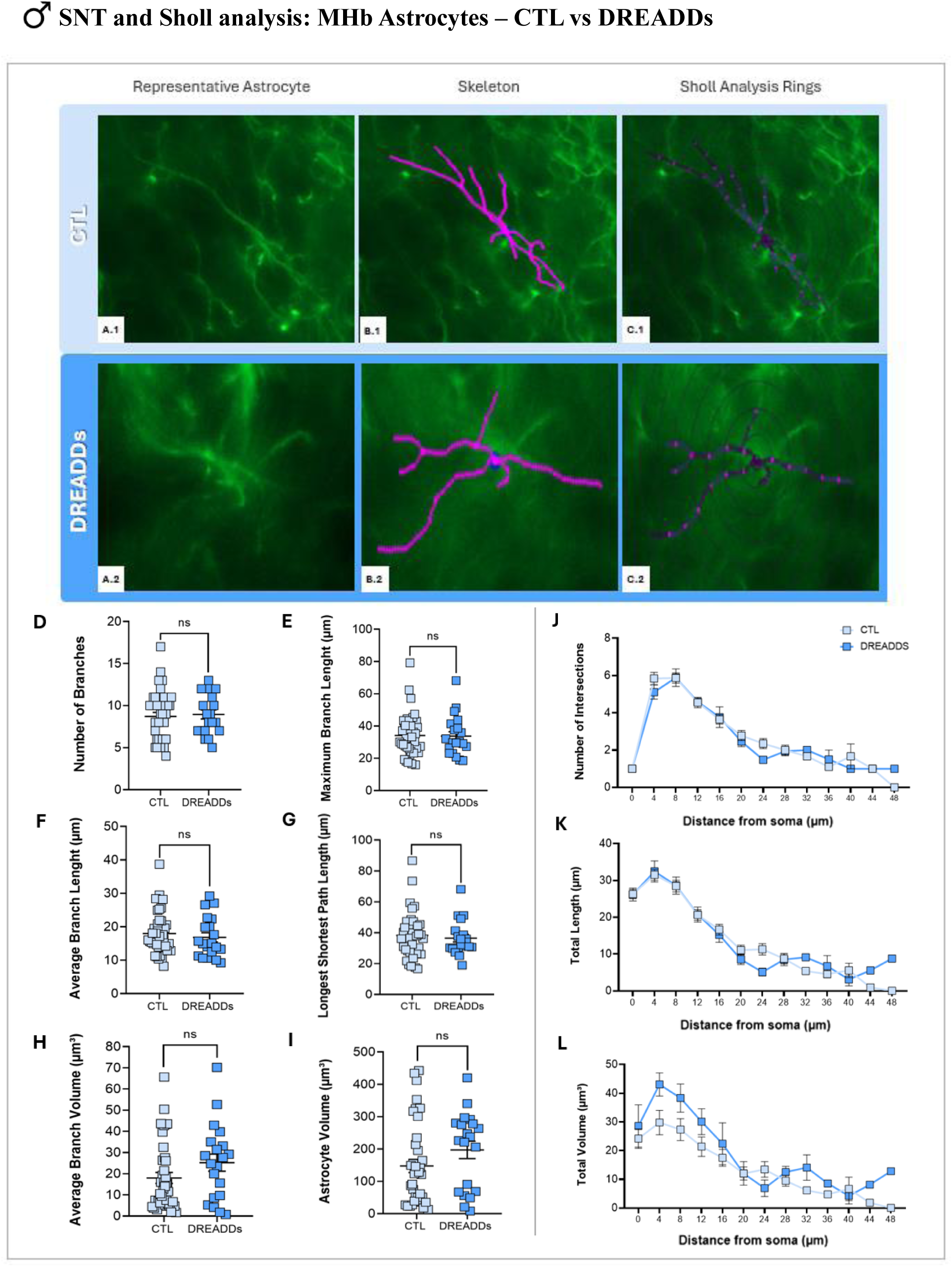
Astrocyte morphological analysis via SNT and Sholl tools under chemogenetic manipulation (CTL vs DREADDs) in the MHb of male mice. **A-C**: Representative images for control group (CTL: A.1-C.1) and experimental group (DREADDs: A.2-C.2). Magnified representative astrocyte (A; scale bar, 10 µm), semi-automated reconstruction skeleton using Simple Neurite Tracer (SNT) (B), and corresponding concentric rings for Sholl analysis (C). **D–I**: SNT-derived parameters quantified per astrocyte. **D**: number of branches, **E**: maximum branch length, **F**: average branch length, **G**: longest shortest path length, **H**: average branch volume, and **I**: total astrocyte volume. **J–L**: Sholl analysis profiles quantified as a function of distance from the soma (4 µm intervals). **J**: number of intersections, **K**: total branch length, and **L**: total branch volume. A two-tailed unpaired Student’s T-test was used for comparisons. Normal distribution and equal variances were assumed. Values are expressed as mean *±* SEM. Significance levels are indicated as follows: ns = not significant. *Note: all animals in this cohort underwent a fear conditioning protocol, including the administration of CNO in the acquisition phase*.

In contrast, Figure 14 reveals a clear, opposing pattern in the female cohort, where DREADDs activation reduced overall astrocyte volume. SNT parameters measured per cell showed a statistically significant decrease in both the average branch volume (Figure 14, H; *p < 0.05) and the total astrocyte volume (Figure 14, I; *p < 0.05). This reduction in volume is further supported by the Sholl analysis graphs (Figure 14, L), where the DREADDs curves sit visibly below the CTL group, showing a decrease in branch volume distribution across most concentric intervals. No significant changes were observed in branch numbers or lengths (Figure 14, D-G).

**Figure 14.**
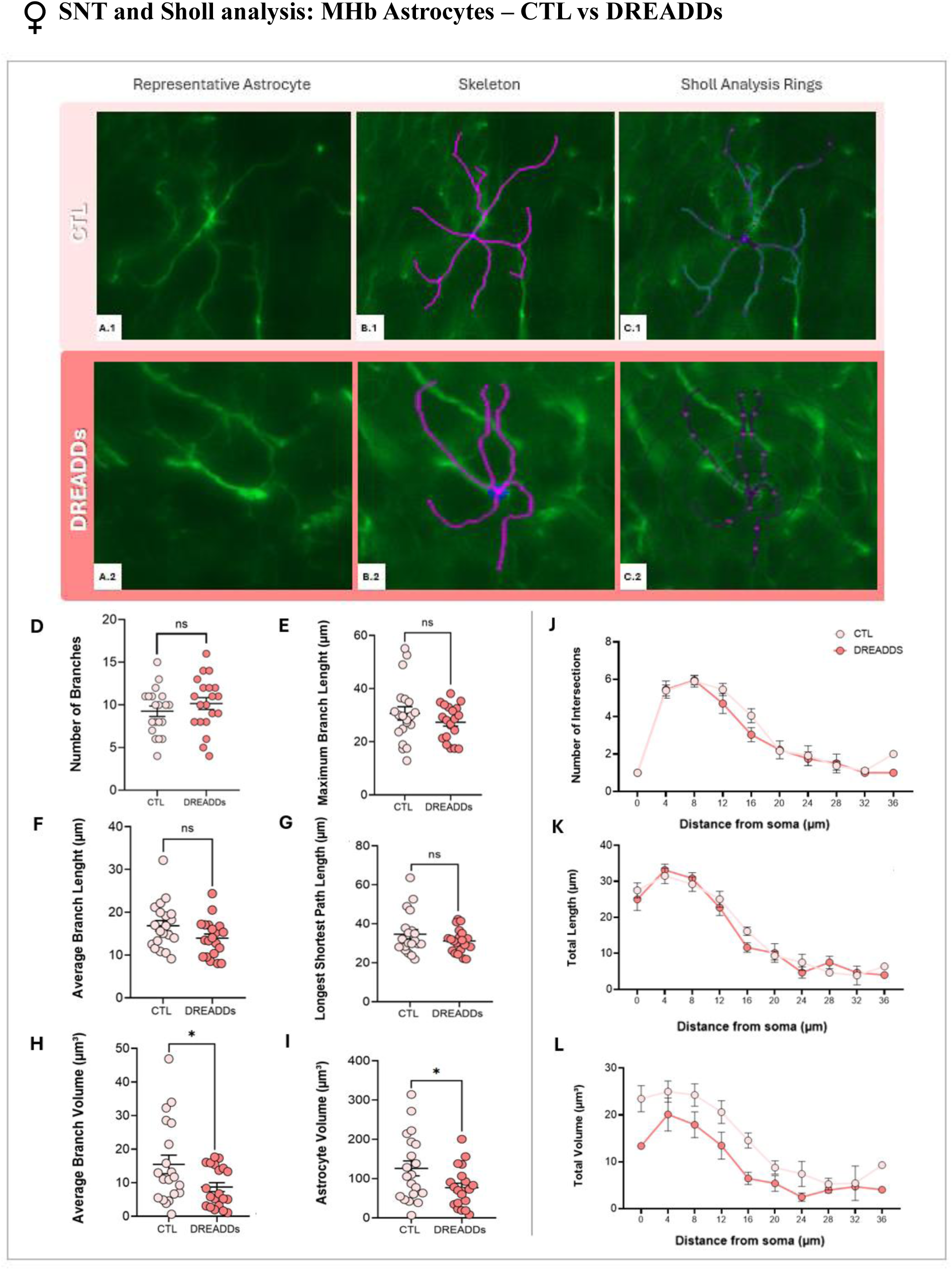
Astrocyte morphological analysis via SNT and Sholl tools under chemogenetic manipulation (CTL vs DREADDs) in the MHb of female mice. **A-C**: Representative images for control group (CTL: A.1-C.1) and experimental group (DREADDs: A.2-C.2). Magnified representative astrocyte (A; scale bar, 10 µm), semi-automated reconstruction skeleton using Simple Neurite Tracer (SNT) (B), and corresponding concentric rings for Sholl analysis (C). **D–I**: SNT-derived parameters quantified per astrocyte. **D**: number of branches, **E**: maximum branch length, **F**: average branch length, **G**: longest shortest path length, **H**: average branch volume, and **I**: total astrocyte volume. **J–L**: Sholl analysis profiles quantified as a function of distance from the soma (4 µm intervals). **J**: number of intersections, **K**: total branch length, and **L**: total branch volume. A two-tailed unpaired Student’s T-test was used for comparisons. Normal distribution and equal variances were assumed. Values are expressed as mean *±* SEM. Significance levels are indicated as follows: ns = not significant, *p < 0.05*. Note: all animals in this cohort underwent a fear conditioning protocol, including the administration of CNO in the acquisition phase*.

## 4. DISCUSSION

This study provides an initial morphological characterization of astrocytes in the MHb of male and female mice across three experimental conditions: metabolic stress (HFD), systemic immune challenge (LPS), and chemogenetic manipulation of astrocyte activity (Gi - DREADDs). Despite the small size of the MHb and its understudied status relative to the LHb (5, 7), its astrocyte population underwent clear morphological remodeling under all three conditions, with responses that differed not only between groups but also consistently between sexes. Combining a 2D skeleton-based approach with SNT, Sholl analysis, and S100β-based density quantification allowed us to capture these changes from complementary perspectives. The goal of this work is to establish a potential glial link between well-known treatments that cause mood disturbances (e.g., HFD, LPS) and the central nervous system (e.g., MHb) in males and females.

In the HFD model, metabolic stress increased the skeletal complexity of MHb astrocytes selectively in males, which showed more branches (Figure 3), whereas females showed no significant change (Figure 4). The male response agrees with prior reports linking HFD to astrocyte structural remodeling (13, 14). Conversely, the absence of a response in female mice is consistent with the view that the central glial response to a HFD depends on both sex and timing: males gain weight and fat before females and therefore tend to show earlier changes in brain glial cells (20). At the 12-week time point examined here, female MHb astrocytes may not yet have reached the remodeling threshold already evident in males, suggesting that a longer exposure would be required to elicit a comparable response. It is worth noting that, in that same study, the response of hypothalamic astrocytes to a HFD was more pronounced in females and differed from what we observed in the MHb (20), a contrast which, once again, points to region-specific astrocyte behavior rather than a uniform effect of the diet

The immune challenge produced the most pronounced sex difference. LPS subtly reduced the complexity of MHb astrocytes in males (fewer branches; Figure 5) and markedly in females, who showed significant reductions in the number of branches, the length of the longest–shortest path, and the maximum branch length (Figure 6). The female’s sensitivity in inflammation-induced astrocytic responses has already been reported in other mood-related regions (21,22).

This was reinforced by the quantification of S100β: males showed no significant changes (Figure 7), whilst LPS-treated females showed a significant decrease in the number of S100β+ astrocytes detected, together with a non-significant trend towards a larger cell area (Figure 8). Although this type of morphology is frequently described in reactive astrocytes, we cannot classify these cells as reactive based solely on morphological criteria.

Turning to the central manipulation, the chemogenetic tool increased the skeletal complexity of MHb astrocytes in both sexes, raising the number of branches and the longest-shortest path length in males (Figure 9), and the number of branches and average branch length in females (Figure 10). This is counterintuitive, since Gi- DREADDs is expected to decrease astrocyte signaling. However, the observed increase in morphological complexity suggests a possible adaptation rather than a simple reduction in activity (16).

Since the AAV was delivered into the MHb, hippocampal astrocytes were analyzed as a distal, non-targeted reference region to test whether the morphological effects were specific to the injection site. No significant change was found in CA1 astrocytes of males (Figure 11), whereas females showed a different pattern: a higher number of branches but a shorter average branch length (Figure 12) (in contrast to the increased average branch length seen in the MHb). These regional differences indicate that the morphological effects of the manipulation are specific to the MHb rather than a generic astrocytic response across the entire brain. This is consistent with the well-established regional heterogeneity of astrocytes (17–19), which are molecularly, functionally, and morphologically distinct across brain regions.

Within the DREADDs group, SNT and Sholl analysis (applied in the MHb) added information that the skeleton method could not capture (unlike the skeleton analysis, which is performed on a 2D maximum-intensity projection, SNT was run on the full image stack and yielded volumetric measurements). Whereas skeletonization indicated increased complexity in both sexes (Figures 9 and 10), the volumetric reconstructions exposed a sex divergence in astrocyte volume: males trended toward increased branch and total cell volume (Figure 13), while females showed a significant reduction in both (Figure 14). The apparent contradiction in females (more and longer branches in 2D, but less volume in the SNT reconstruction) suggests that they do not hypertrophy under chemogenetic manipulation but rather thin and redistribute their processes, becoming more ramified while occupying a smaller territorial volume. This reinforces the value of complementing skeletonization with volumetric reconstruction.

Taken together, although the direction of the remodeling differed between challenges (a gain in complexity under HFD and DREADDs, and a loss under LPS), the three models converge on the same conclusion: astrocytes in the MHb are morphologically sensitive, and this response is consistently sexually dimorphic.

Several limitations constrain these initial findings. The most important issue is the relatively small sample size of some cohorts (particularly the LPS group), which limits statistical power. The in-depth analyses are also unevenly distributed: SNT and Sholl analyses were applied only to the DREADDs MHb, and S100β density was assessed only in the LPS group; both are currently being extended to the remaining groups and to both sexes. In addition, the hippocampus was the only reference region examined, when a comparison with additional emotion-related regions would be informative. It is important to emphasize that morphology alone cannot establish whether an astrocyte is reactive; reactivity must be assessed using multiple molecular and functional parameters (23); therefore, our morphological data must be complemented by transcriptomic approaches to identify markers associated with reactivity, such as complement component C3 (12), and by the profile of pro-inflammatory cytokines, before a functional status can be assigned to the structural changes described. The markers used in immunofluorescence also have some limitations: GFAP mainly labels the cytoskeleton and only a subset of the astrocyte population, thereby underestimating their fine processes and true extent, whilst S100β is less specific to astrocytes than GFAP, as it is also expressed in oligodendrocytes and some neurons (24). The strategy of using GFAP for morphology and S100β for density is reasonable but would be strengthened by including additional astrocytic markers such as Sox9 (25). Finally, the DREADDs cohort underwent fear conditioning prior to perfusion, and stress itself remodels astrocytes (22); therefore, some of the sex differences relating to baseline and to DREADDs may reflect sex-specific responses to that stress.

In addition to carrying out these analyses across all groups and both sexes, the next crucial step will be to correlate the specific morphological characteristics described here with the behavioral data already collected from these cohorts. This integration would reveal whether the various structural changes observed in MHb astrocytes are directly linked to sex-specific emotional phenotypes.

## 5. CONCLUSIONS

In summary, this study shows that MHb astrocytes undergo sex-dependent morphological remodeling in response to metabolic, immune and chemogenetic challenges. Chronic exposure to a HFD increased astrocytic complexity selectively in males, whereas female MHb astrocytes showed no significant structural changes over this time course. Systemic inflammation, by contrast, tended to reduce MHb astrocytic complexity, particularly in females, who also exhibited a lower density of S100β+ cells. Chemogenetic manipulation triggered sex-specific changes as well: branching complexity increased in both sexes, while volumetric adaptations diverged between males and females. These responses appear to be region-specific, as hippocampal astrocytes displayed distinct remodeling patterns (at least in the DREADDs group). Taken together, these findings point to a compelling sexual dimorphism in how MHb astrocytes respond to peripheral and central challenges, although further studies with larger cohorts will be required to overcome current sample-size limitations.

## ACKNOWLEDGEMENTS

The authors would like to thank the Animal Facility and the Microscopy Unit of the General Research Services (SGIker, UPV/EHU) for their technical and human support.

